# Rod phototransduction and light signal transmission during type 2 diabetes

**DOI:** 10.1101/2020.01.14.905265

**Authors:** Silke Becker, Lara S. Carroll, Frans Vinberg

**Author notes:** Corresponding Author: Silke Becker PhD, John A. Moran Eye Center, 65 Mario Capecchi Drive, University of Utah, Salt Lake City, Utah, USA. Email addresses of co-authors.

## Abstract

**Objective:** Diabetic retinopathy is a major complication of diabetes recently associated with compromised photoreceptor function. Multiple stressors in diabetes, such as hyperglycemia, oxidative stress and inflammatory factors, have been identified, but systemic effects of diabetes on outer retina function are incompletely understood. We assessed photoreceptor physiology *in vivo* and in isolated retinas to better understand how alterations in the cellular environment compared to intrinsic cellular/molecular properties of the photoreceptors, affect light signal transduction and transmission in the retina in chronic type 2 diabetes.

**Research Design and Methods:** Photoreceptor function was assessed in BKS.Cs-Dock7^m^ +/+ Lepr ^db^/J mice, using homozygotes for Lepr^db^ as a model of type 2 diabetes and heterozygotes as non-diabetic controls. *In vivo* electroretinogram (ERG) was recorded in dark adapted mice at both 3 and 6 months of age. For *ex vivo* ERG, isolated retinas were superfused with oxygenated Ames’ media supplemented with 30 mM glucose or mannitol as iso-osmotic control and electrical responses to light stimuli were recorded.

**Results:** We found that both transduction and transmission of light signals by rod photoreceptors were compromised in 6 month old (n=9-10 eyes from 5 animals,*** p<001) but not in 3 month old diabetic mice *in vivo* (n=4-8 eyes from 2-4 animals). In contrast, rod signaling was similar in isolated retinas from 6 month old control and diabetic mice under normoglycemic conditions (n=11). Acutely elevated glucose *ex vivo* increased light-evoked rod photoreceptor responses in control mice (n=11, *** p<0.001), but did not affect light responses in diabetic mice (n=11).

**Conclusions:** Our data suggest that long-term diabetes does not irreversibly change the ability of rod photoreceptors to transduce and mediate light signals. However, type 2 diabetes appears to induce adaptational changes in the rods that render them less sensitive to increased availability of glucose.

**Significance of this study:** *What is already known about this subject?:* Retinal neuronal dysfunction in diabetic retinopathy frequently precedes microvascular changes. Damage to photoreceptors has been reported, although the literature is not entirely conclusive, and the underlying mechanisms and reversibility of photoreceptor dysfunction in diabetes remain unknown.

*What are the new findings?:* We confirm that retinal function is impaired *in vivo* in diabetic mice, but report normal function of the isolated diabetic retina under optimal physiological conditions. However, elevated glucose concentrations augment photoreceptor function in the isolated non-diabetic retina, whereas this response is impaired by long-term diabetes.

*How might these results change the focus of research or clinical practice?:* Our results clarify that photoreceptor function is not irreversibly damaged and that systemic factors may suppress photoreceptor function in diabetic mice *in vivo*. The consequences of altered glucose handling in long-term diabetic photoreceptors remain to be evaluated.

## INTRODUCTION

Diabetic retinopathy, one of the leading causes of irreversible vision loss worldwide ^1^, has traditionally been characterized by retinal microvascular abnormalities. Recent studies in human patients with diabetes suggest, however, that retinal diabetic neuropathy with neuroglial dysfunction and degeneration frequently precedes visible microvasculopathy ^2 3^. Even in the absence of, or with minimal signs of retinal vascular abnormalities, patients with diabetes show dysfunction and loss of retinal neuronal cells. In fact, impairment of the neurovascular unit has been suggested to lead to or accelerate retinal vascular disease progression, since implicit time delay of the multifocal electroretinogram (ERG) predicts onset of visible vascular changes ^4^. Damage to nerve fiber, ganglion cell and inner plexiform layers in the human eye has been demonstrated in diabetes by impaired pattern ERG and oscillatory potentials ^5^ and thinning of inner retinal layers ^3^. Reduced amplitudes and longer implicit times of the scotopic and photopic b- waves ^5^ also implicate dysfunction of rod and cone bipolar cells in retinal diabetic neuropathy of human patients. These findings are not universal, however, and reduced scotopic a- and b-wave amplitudes have not been shown in all studies ^6^.

Murine models of diabetes share many features of retinal dysfunction and cell loss with patients with diabetes. The positive component of the scotopic threshold response, a measure of retinal ganglion cell (RGC) function, is attenuated following streptozotocin (STZ)-induced type 1 diabetes in the rat ^7^. Additionally, thickness of the nerve fiber and RGC layer, and RGC density are all reduced in STZ-induced rodents ^3^. Loss or reduced inhibitory output of amacrine cells has been observed in type 1 diabetes models ^8-12^, and may underlie reduced amplitude and delayed implicit time of oscillatory potentials ^7 13^. Similarly to human studies, outer retinal dysfunction and cell loss in diabetic animal models remain controversial. Rod photoreceptor and bipolar cell dysfunction, based on reduced amplitudes and delayed implicit times of the scotopic a- and b-waves, were reported in the db/db mouse, a type 2 diabetes model ^14^. Photoreceptor death was also reported in a rat STZ model ^15^, but was not validated in other studies ^16 17^ which reported no or only minor changes in scotopic ERG ^7^, or even increased photoreceptor function in Zucker Diabetic Fatty (ZDF) rats, another type 2 diabetes model ^18^.

Several studies have implicated photoreceptors in the pathogenesis of diabetic retinopathy ^19^. Rod photoreceptors are a significant source of oxidative stress in diabetic animals ^17^ and rod degeneration or blocking of visual transduction protects from vascular diabetic retinopathy ^20-24^. However, systemic or cell-intrinsic mechanisms affecting photoreceptor function during diabetes are not well understood. Transient hyperglycemia in patients with diabetes has been associated with increased scotopic b- and photopic a- and b-wave amplitudes ^25^. While the multifocal ERG waveform is delayed in normoglycemic patients with diabetes compared to healthy volunteers, this is partly reversed by elevation of blood glucose to their usual level of (habitual) hyperglycemia ^26 27^. These findings indicate that both acute hyperglycemia and chronic diabetes modulate retinal signaling. We provide evidence for our hypothesis that long-term uncontrolled type 2 diabetes in db/db mice does not permanently alter intrinsic mechanisms in rod photoreceptors responsible for transducing and transmitting light signals. However, we find that rod photoreceptors in the *ex vivo* isolated diabetic retina are less sensitive to acute elevation of extracellular glucose. We suggest that attenuation of the *in vivo* ERG in diabetic mice is likely due to diabetes-induced systemic factors influencing rod photoreceptors, which once removed, can no longer mediate suppression of photoreceptor responses in the context of the *ex vivo* testing.

## RESEARCH DESIGN AND METHODS

### Animals

Male and female C57B/6J mice and BKS.Cg-Dock7^m^ +/+ Lepr^db^/J mice were bred in house or purchased at 8-10 weeks of age from Jackson Laboratory. Animals homozygous for the Lepr^db^ mutation (db/db) are diabetic. Age-matched heterozygotes (db/+) were used as non-diabetic controls.

### Weight, blood glucose and HbA1c

Animals were kept under a 12:12 hour light/dark cycle and given ad libitum access to food and water. Diabetic status was confirmed by measuring weight, blood glucose and HbA1c at 3 or 6 months of age (Supplemental Figure 1).

Blood glucose concentrations during isoflurane anesthesia were assessed in C57B/6J mice immediately before anesthesia and 20 min after beginning inhalation of 2% isoflurane.

### *In vivo* ERG

Mice were fully dark adapted and anesthetized with 1.8-3.0% isoflurane in room air using a SomnoSuite Low Flow Anesthesia System (Kent Scientific). Pupils were dilated with 2% atropine sulfate eye drops (Akorn, Illinois) and corneas lubricated with a mixture of phosphate buffered saline (PBS) with Goniovisc (2.5% hypromellose solution, Contacare, Gujarat, India). Body temperature was maintained by placement on a heated table throughout the experiment. ERG waveforms were recorded using corneal contact lens electrodes and subdermal needle electrodes between the eyes as reference and at the base of the tail as ground electrodes. Responses to light flashes (rod stimuli, to activate rod phototransduction in rodents), were collected using a ColorDome and Espion V6 software (Diagnosys LLC).

For scotopic ERGs 20 sweeps with inter-sweep intervals of 2 s were recorded at light flash energies (Luminous energy, Q_V_) from 10^−5^ to 10^−3^ cd.s.m^-2^, 10 sweeps with 5 s intervals at 10^−2^ cd.s.m^-2^, 5 sweeps with 10 s intervals from 10^−1^ to 0.5 cd.s.m^-2^, 3 sweeps with 20 s intervals from 1 to 5 cd.s.m^-2^, and 2 sweeps with a 60 s interval at 10 and 30 cd.s.m^-2^. Oscillatory potentials were high-pass filtered above 25 Hz and subtracted from ERG waveforms. a- and b-wave response amplitudes were measured between baseline and trough and between trough and peak respectively, and were plotted as the response amplitude over the log light intensity. a- and b-wave implicit times were determined as time from stimulus onset to peak. The amplification constant *A* of phototransduction at 1 cd.s.m^-2^ was calculated as described by Lamb & Pugh ^28^.

### *Ex vivo* ERG

Mice were fully dark adapted overnight and sacrificed by CO_2_ asphyxia. Eyes were enucleated and dissected in Ames’ media supplemented with 0.25 mM sodium glutamate ^29^ and oxygenated with 95% O_2_/5% CO_2_. The anterior eye and the lens were removed and retinas were carefully dissected and mounted on an *ex vivo* specimen holder, as previously described ^30^. Tissue was superfused with oxygenated Ames’ media containing 0.25 mM sodium glutamate heated to 36**°**C. Dissected eye cups from fellow eyes were stored in oxygenated Ames’ media at room temperature for up to 2 hours. ERG waveforms were elicited by light stimuli of increasing intensity at 530 nm, amplified using a DP-311 differential amplifier (Warner Systems), digitized with a MEA2100-System (Multichannel Systems), and recorded using Multi Channel Experimenter software. To investigate acute effects of high glucose, we added either 30 mM D-glucose or mannitol as iso-osmotic control to the perfusate in fellow eyes 20 min before recordings were started.

To determine ON-bipolar cell (BPC, also termed PII) responses, combined photoreceptor (PR, also termed fast PIII) and BPC responses were measured at light intensities from 3 to 21 photons/μm^2^ in the presence of 100 μM BaCl_2_ (to inhibit K^+^ channels in Müller glia) ^30^. Corresponding PR responses were recorded in the presence of 100 μM BaCl_2_ and 40μM DL-AP4 (to block neurotransmission from photoreceptors to ON-bipolar cells) and subtracted from combined PR+BPC responses to obtain PII responses ^31^ (Figure 2A-C). PR response families were recorded at light intensities from 33 to 1,045 photons μm^-2^ in the presence of 100 μM BaCl_2_ and 40 μM DL-AP4. The amplification constant *A* for phototransduction at a light intensity of 50 photons/μm^2^ was calculated, as previously described ^28^.

**Figure 1:**
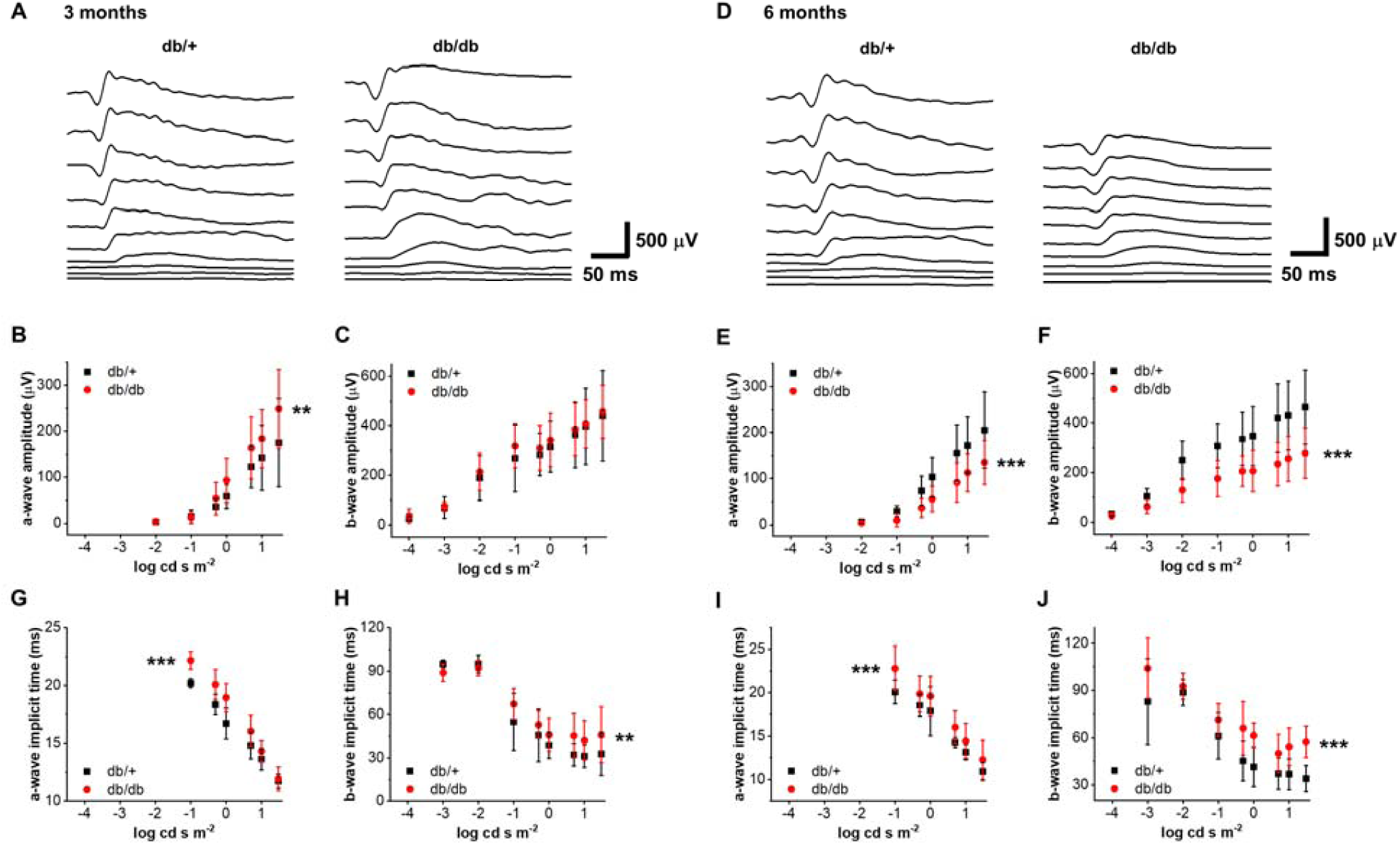
Scotopic a- and b-waves are reduced in 6 but not 3 months old type 2 diabetic mice *in vivo*. Representative *in vivo* ERG light responses from 3 (A) and 6 (D) month old control (db/+) and diabetic (db/db) mice to flashes of light ranging from 10^−5^ to 30 cd s m^-2^. The scotopic a-wave (B, E) and b-wave (C, F) amplitudes as a function of flash energy (Q_V_) for db/+ (black squares) and db/db (red circles) mice at 3 (A-C, n=4-8 eyes from 2-4 animals, ** p<0.01) and 6 (D-F, n=9-10 eyes from 5 animals, *** p<0.001) months of age. a- and b-wave implicit times in db/db (black squares) mice compared to db/+ (red circles) mice at either time point (G-J, ** p<0.01, *** p<0.001).

**Figure 2:**
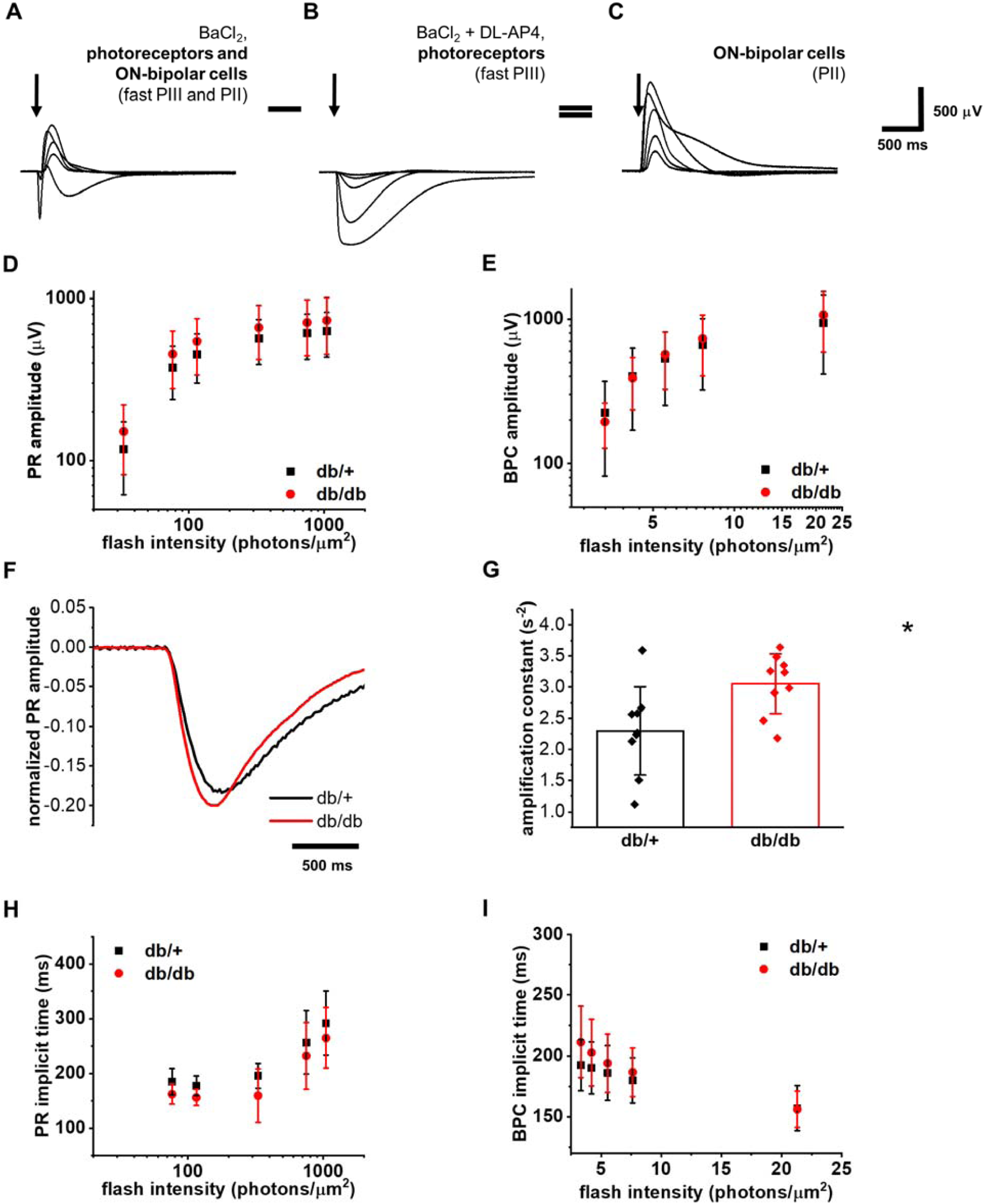
Isolation of photoreceptor and ON-bipolar cell function by *ex vivo* ERG demonstrates no amplitude changes in the diabetic retina. The PR response family as a measure of *ex vivo* photoreceptor function in this example trace was recorded in the presence of BaCl_2_ and DL-AP4 with light stimuli (arrows, B). Combined PR and BPC responses were obtained at the same light intensities in the presence of BaCl_2_ alone (A) and isolated ON-bipolar cell responses were calculated by subtraction of PR from combined PR+BPC waveforms (C). PR (D) and BPC (E) amplitude data for db/+ (black squares) and db/db (red circles) mice in normoglycemic conditions (D-E, n=11 for PR and n=4-6 for BPC). Averaged PR waveforms at 50 photons μm^-2^ normalized to maximal response amplitude at 550 photons μm^-2^ (F), and quantification of the amplification constant (G, n=9, * p<0.05, see text for details). PR (H, n=11, * p<0.05) and BPC (I, n=4-6) implicit times in db/db (red circles) compared to db/+ (black squares) mice.

### Statistical analysis

Data are displayed as arithmetic means ± standard deviation. Statistical analysis was performed using Origin software. Differences between two groups with independent values were determined by two-tailed student’s t-test, for multiple stimulus intensities by two-way ANOVA.

### Data and Resource Availability

No datasets and no applicable resources were generated or analyzed during the current study.

## RESULTS

### Type 2 diabetes has different effects on outer retinal physiology under *in vivo* and *ex vivo* conditions

Long-term diabetes impairs the function and survival of RGCs and amacrine cells ^3 5 7-9^, but there is currently no consensus on its effects on phototransduction and synaptic transmission of the rod photoreceptors. Here we first asked: what is the time course of alterations in rod-mediated light signals caused by type 2 diabetes in habitual blood glucose? To answer this question, we compared scotopic *in vivo* ERG a- and b-waves in db/db mice and their age-matched db/+ controls at 3 and 6 months of age. We used isoflurane anesthesia, which in contrast to ketamine/xylazine anesthesia, did not significantly increase blood glucose in C57BL/6J mice (Supplemental Figure 1C, ^32 33^).

We did not observe reduced *in vivo* a- and b-wave amplitudes in diabetic compared to non-diabetic control mice at 3 months of age, but rather a slight increase of the *in vivo* a-wave amplitude, similar to that reported by Johnson *et al*. ^18^ (Figure 1A-C). a- and b-wave implicit times were delayed at this age, suggesting beginning photoreceptor and ON-bipolar cell dysfunction (Figure 1G-H). Conversely, in 6 month old diabetic mice the scotopic *in vivo* ERG a- and b-wave amplitudes were reduced (Figure 1D-F) and a- and b-wave implicit times delayed (Figure 1I-J). The amplification constant (*A*) of phototransduction, a measure of signal amplification in photoreceptors, was determined based on the leading edge of the a-wave, when b-wave intrusion is negligible, as previously described ^34^. *A* was not significantly different between 6 month old control and diabetic mice (n=8-9, p=0.25), indicating that the activation reactions of phototransduction in rods are not affected by diabetes. Interestingly, we found a trend towards reduced b/a-wave ratio in diabetic animals compared to non-diabetic controls at 3 months of age, which was highly significant at 6 months of age (Table 1).

**Table 1:** b/a-wave ratios in diabetic mice at 3 and 6 months of age.

| Age | db/+ | db/db | p value |
| --- | --- | --- | --- |
| **3 months** | 3.12 ± 0.73 | 2.76 ± 0.57 | p=0.42 |
| **6 months** | 2.65 ± 0.27 | 2.20 ± 0.23 | ***p<0.001 |

Although *in vivo* ERG data by us and others suggest that the light-sensitive current and/or synaptic transmission of rods are compromised during long-term hyperglycemia, it is not known whether these changes are related to an altered extracellular environment or permanent molecular/structural changes in rods or rod bipolar cells caused by diabetes. Moreover, systemic effects of anesthesia, such as hyperglycemia, complicate interpretation of the results obtained by *in vivo* ERG in animal models of diabetes. To determine the effects of chronic diabetes on the intrinsic functional properties of rods and rod bipolar cells in a normoglycemic environment and to avoid systemic complications of anesthesia, we assessed rod phototransduction and synaptic transmission using *ex vivo* ERG in 6 month old control and diabetic mice. This technique also allowed us to pharmacologically isolate and quantify the rod photoreceptor (PR) and ON-bipolar cell (BPC) components of the ERG signal (Figure 2A-C). Surprisingly, in contrast to the *in vivo* ERG, the amplitudes of the PR or BPC components were not significantly changed in 6 month old db/db mice as compared to age-matched db/+ mice (Figure 2D-E). These results demonstrate that transduction or transmission of light signals in rods is not adversely affected by long-term type 2 diabetes under the standard physiological environment used in *ex vivo* experiments.

Although our amplitude analysis of the *ex vivo* ERG data suggests that diabetes does not alter the light sensitivity of rod photoreceptors in normal glucose, it is possible that the kinetics of phototransduction activation and inactivation reactions are modulated by diabetes. To test this, we determined the gain of the phototransduction activation reactions and time-to-peak (=implicit time) from *ex vivo* ERG PR responses to dim flashes (Figure 2F-G). Interestingly, we found that long-term type 2 diabetes increased *A* and contrary to *in vivo* ERG findings, no change was seen in the *ex vivo* PR or BPC time to peak (Figure 2H-I).

### Acute hyperglycemia promotes light responses in rod photoreceptors of non-diabetic, but not diabetic (type 2) mouse retinas

Previous clinical studies demonstrating increased b-wave amplitudes following elevation of blood glucose suggest that retinal function may be modulated by extracellular glucose concentration ^25^. To investigate this, we used the *ex vivo* ERG method that allows us to precisely control glucose concentrations. We first evaluated whether acute elevation of extracellular glucose in the perfusate altered light evoked activity of the rod photoreceptors or bipolar cells in C57BL6/J retinas. Supplementation with 30 mM glucose compared to mannitol iso-osmotic control increased the maximal PR response amplitude (Figure 3A, C). BPC responses were also slightly increased by acute addition of glucose (Figure 3B, D). However, PR/BPC amplitude ratio was not significantly changed between mannitol and glucose supplementation, indicating that the increased BPC amplitude in the presence of added glucose derives from increased photoreceptor responses.

**Figure 3:**
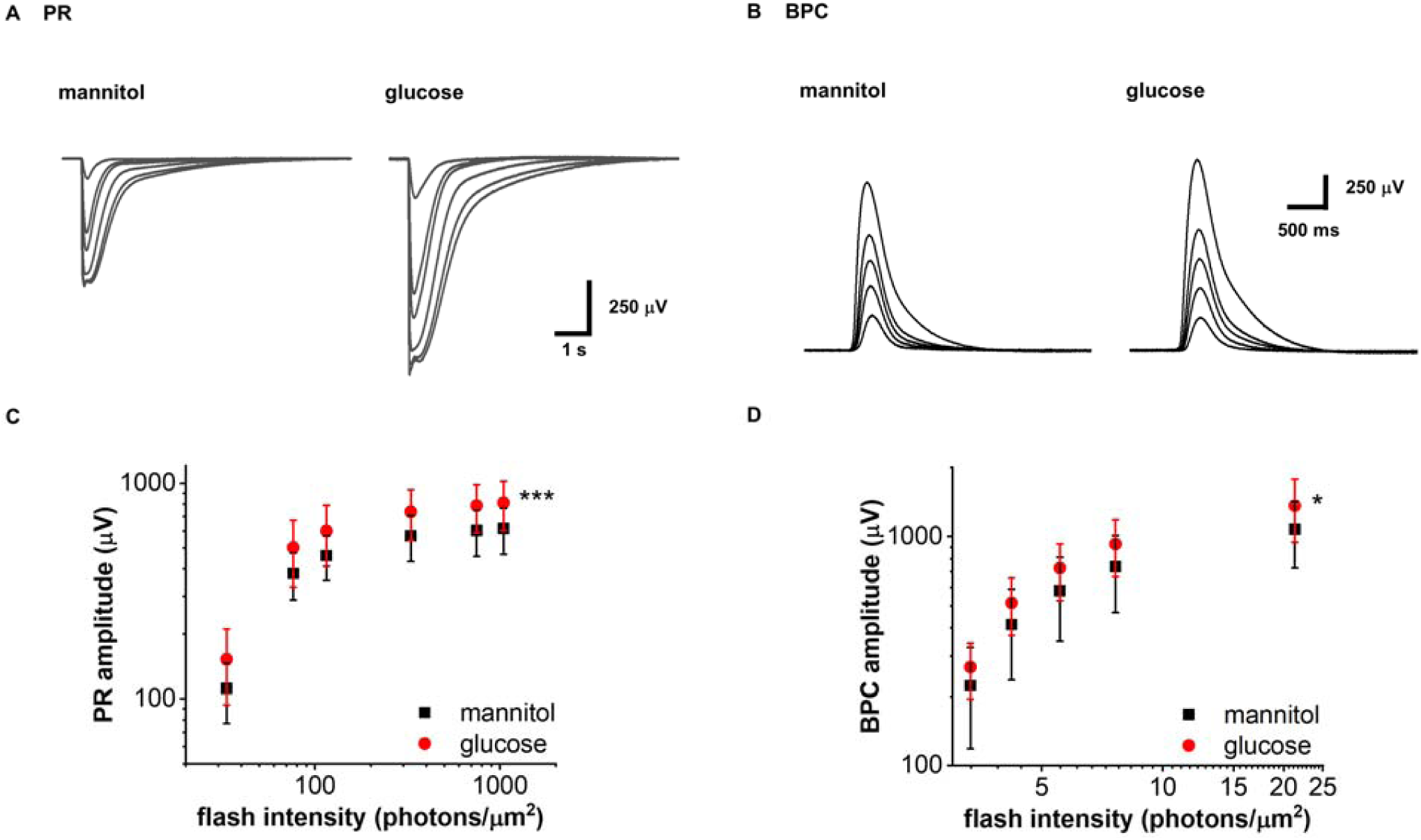
Acute elevation of extracellular glucose increases photoreceptor amplitudes of wild-type mice. Representative *ex vivo* ERG PR (A) and BPC (B) light response families in normal (iso-osmotic mannitol control, normal glucose + 30 mM mannitol) and after supplementation with 30 mM glucose. PR (C, *** p<0.001, n=7) and BPC (D, * p<0.05, n=5) amplitude data in normal (mannitol control, black squares) and high glucose (red circles) in C57BL6/J mice.

Next, we wanted to know if long-term diabetes alters the acute hyperglycemia-induced augmentation of photoreceptor responses. Diabetes has been shown to cause various long-term adaptations in the retina, including differential expression of glucose transporters ^35^. Thus, we hypothesized that the photoreceptors in 6 months old db/db mice are less sensitive to elevated extracellular glucose. Indeed, PR amplitudes of db/+ control retinas were significantly augmented in acute hyperglycemia, whereas the photoreceptor responses of db/db retinas were not affected by high glucose exposure (Figure 4A-C). On the other hand, we did not observe a significant modulation of the BPC amplitudes by elevated glucose exposure in control or diabetic mice (Figure 4D-F). PR implicit times were delayed in high glucose compared to mannitol iso-osmotic control in both db/+ and db/db retinas (Figure 4G, H). While the PR implicit times in db/db retinas were accelerated compared to db/+ retinas in normal glucose (Figure 2H), no significant difference was observed between non-diabetic retinas in *normal glucose* and diabetic retinas in *high glucose* (compare black squares in Figure 4G and red circles in Figure 4H). The amplification constant *A* as described above was not different between high glucose and mannitol treatment in either db/+ or db/db mice (data not shown), suggesting that the increased implicit times of the rod responses must be due to decelerated phototransduction deactivation rather than activation reactions caused by the elevated glucose. Similar to the PR times to peak, the BPC implicit times lengthened by supplementation with glucose compared to mannitol in both non-diabetic and diabetic animals (Figure 4I-J).

**Figure 4:**
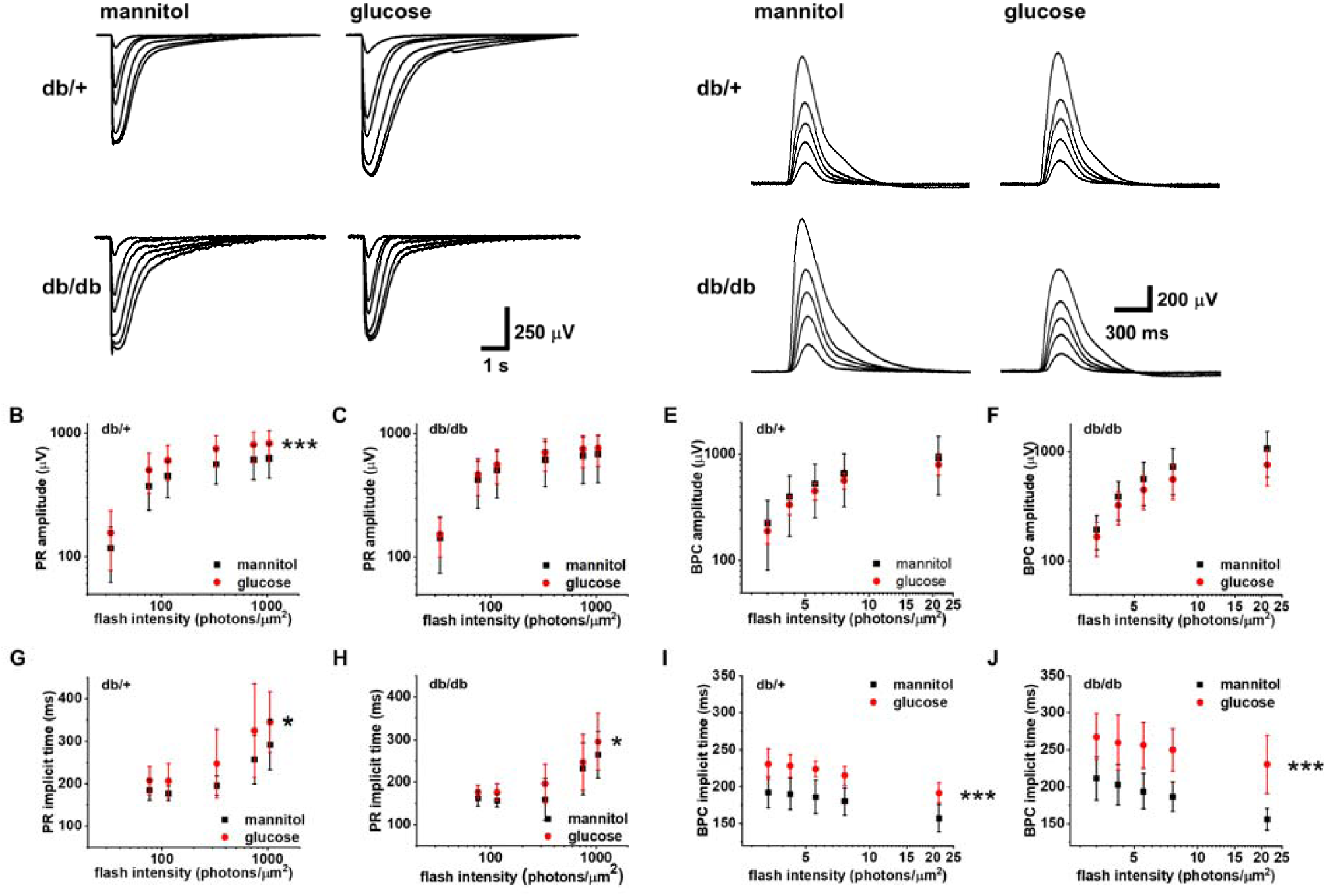
Long-term diabetes modulates the response to acute hyperglycemia in photoreceptors. E*x vivo* ERG PR (A-C, n=11 retinas) and BPC (D-F, n=6 db/+ and n=4 db/db retinas) light response families and amplitude data in normal (iso-osmotic mannitol control, normal glucose + 30 mM mannitol, black squares) and after supplementation with 30 mM glucose (red circles) in 6 month old control (db/+, *** p<0.001, B, E) and diabetic (db/db, C, F) mice. PR (G, H) and BPC (I, J) implicit times in normal (black squares) and high (red circles) glucose in db/+ (G, I) and db/db (H, J) mice (G-H, * p<0.05, I-J, *** p<0.001).

## CONCLUSIONS

It is widely accepted that diabetes results in loss of RGC and amacrine cells and associated attenuation of oscillatory potentials, both in diabetic animal models and patients ^3 7 8 36^. Effects of diabetes on photoreceptor and ON-bipolar cell light signaling remain controversial, however. While many studies demonstrated attenuated a- and b-wave amplitudes of the scotopic *in vivo* ERG ^9 14 37 38^, other reports are contradictory ^7 18 39^. Causes of these discrepancies remain unclear, but have previously been ascribed to duration of diabetes ^9 37^. This is supported by our study, which shows reduction of *in vivo* a- and b-wave amplitudes in db/db mice at 6 months of age, whereas the scotopic a-wave was increased in 3 months old diabetic mice (Figure 1 D-F and 1 A-C, respectively). We conclude that changes in *in vivo* rod photoreceptor function are minor at 3 months of age, but significant dysfunction occurs in uncontrolled type 2 diabetes at 6 months of age. Our data also indicate that signal transmission from rods to rod bipolar cells, measured by the b/a-wave ratio, is impaired in db/db mice, which progresses from 3 to 6 months of age.

Importantly, though, our data imply that altered photoreceptor function *in vivo* (Figure 1) may not be explained by intrinsic dysfunction of the light signal transduction or transmission in diabetic rods (Figure 2), but by systemic diabetes-mediated changes that alter rod function. This is consistent with lack of photoreceptor loss ^19^ and absence of reports of reduced scotopic ERGs in patients with diabetes. Since other studies have shown attenuated photoreceptor function in *ex vivo* transretinal recordings from STZ mice ^40^, our findings may be model dependent, or restricted to the type 2 diabetic db/db mouse. It is also possible that compromised rod photoreceptor function in the STZ mouse model derives from neurotoxic effects of STZ ^41^ rather than a consequence of diabetes. Our results are consistent with other *ex vivo* studies showing reduced inhibitory output from amacrine cells but no changes in excitatory output of rods and increased or unaltered ON bipolar cell output in diabetic retinas ^*10 11 42*^.

The *ex vivo* ERG technique uniquely allows precise control of the experimental conditions, including extracellular glucose concentration, and isolation of the ERG waveforms from individual retinal cell types. We therefore conclusively show that under normoglycemic conditions which lack detrimental factors associated with diabetes, such as oxidative stress, generation of light signals and their transmission to rod bipolar cells are not altered by long-term diabetes. What could explain the difference between *in vivo* and *ex vivo* ERG in diabetic mice? One obvious difference is the hyperglycemic environment of photoreceptors in diabetic mice *in vivo* ^43^ compared to normal extracellular glucose in the *ex vivo* experiments for retinas from both control and diabetic mice. However, increasing the glucose concentration in *ex vivo* experiments with diabetic mice did not affect their photoreceptor response amplitudes (Figure 4A, C). Thus, the difference in the extracellular glucose *in vivo* cannot explain the decreased *in vivo* ERG amplitudes.

Other confounding factors include anesthesia-induced systemic effects. Indeed, anesthetic agents are known to affect ERG amplitudes ^44^. Thus, one possibility is a different response of retinas in diabetic mice to the systemic changes caused by anesthesia. This is supported by the observation that scotopic a-waves are not altered in conscious human patients with diabetes during *in vivo* ERG experiments ^45^. One obvious systemic effect of anesthesia is elevated blood glucose ^33^. However, previous studies have demonstrated that hyperglycemia in fact promotes retinal signaling in diabetic patients ^25 26^, whereas light-evoked *ex vivo* responses from diabetic mouse retinas were unchanged by hyperglycemia in our study. Thus, it is improbable that anesthesia-induced hyperglycemia could explain reduced *in vivo* ERG a- or b-waves in 6 month old db/db mice. Furthermore, we used isoflurane anesthesia which did not cause anesthesia-induced hyperglycemia. While previous studies concluded that retinal neurodegeneration in db/db mice was abrogated by dietary restriction, it remains unclear whether this was the result of reduced hyperglycemia, body weight or other diabetes-related changes ^14^. Consequently, we suggest that systemic effects, other than hyperglycemia, that are related to diabetes or general anesthesia affect photoreceptor light responses, resulting in reduced a- and b-wave amplitudes *in vivo* after long-term uncontrolled diabetes in mice.

It is also possible that altered recording geometry due to different size or body lipid composition of diabetic animals affects *in vivo* ERG amplitudes. This, however, is not a probable explanation since altered *in vivo* ERG a- and b-waves have been reported both in type 1 (non-obese) and 2 (obese) diabetic animal models. Overall, we conclude that photoreceptor and ON-bipolar cells in diabetic mouse retinas have the potential to mediate normal light signals, although they may have been negatively impacted by altered extracellular environment in diabetes *in vivo*.

While photoreceptors are highly metabolically active and mostly depend on glucose as their source of energy ^46^, effects of acute hyperglycemia on retinal function remain unclear, although previous reports suggest a correlation between bipolar cell function and blood glucose levels. In clinical studies the scotopic b-wave amplitude was increased in hyperglycemic diabetes patients ^25^. Similarly, the scotopic a- and b-waves were reported to be increased in diabetic ZDF rats, while acute insulin treatment resulted in decreased a-wave amplitudes ^18^. These results seem at odds with our observation that acutely elevated glucose increased photoreceptor responses in control but not in 6 months old db/db mice. However, in human patients, glucose levels are typically controlled by insulin, suggesting that the ability to promote photoreceptor responses is lost specifically due to a long-term exposure to an uncontrolled hyperglycemic environment. The difference of our data in comparison to that of Johnson *et al*. remain unclear, but may be related to different animal models and longer duration of diabetes in our study.

While our experiments did not identify the mechanisms by which light-evoked photoreceptor responses are increased in the presence of additional glucose, likely targets include energy dependent processes in photoreceptors such as Na^+^/K^+^ ATPases, Na^+^/Ca^2+^/K^+^ exchangers and Ca^2+^ ATPases ^46^, ATP-sensitive K^+^ channels expressed in photoreceptors ^47^ or other ATP-dependent processes. The main glucose transporters expressed in photoreceptors are the insulin-independent GLUT-1 and insulin-dependent GLUT-4 ^46^, which have previously been reported to be reduced in diabetes ^35 48 49^, thus potentially explaining the inability of the photoreceptors in db/db mice to utilize increased glucose concentrations to promote their light responses. However, these findings are not universal and increased expression of GLUT-1 in diabetes has also been reported ^43^. Determining underlying causes for the lack of increased photoreceptor function during acute hyperglycemia in diabetic animals was beyond the scope of this study and should be subject to future investigations.

In summary, we demonstrate that the rods have potential to transduce and transmit light signals normally in db/db mice at least up to 6 month of age under standard *ex vivo* environment. Differences between diabetic and non-diabetic mice in the *in vivo* ERG are likely due to systemic factors, which remain to be identified.

## Author Contributions

S.B. designed and conducted experiments and wrote the manuscript. L.C. contributed to data interpretation and manuscript preparation, F.V. contributed conceptually to research, designed experiments and wrote the manuscript. S.B. is the guarantor of this work, had full access to the data and takes responsibility for the integrity and accuracy of the data and its analysis.

## Prior Presentation Information

Parts of this study were presented in abstract form at the Association for Research in Vision and Ophthalmology Annual Meeting in Vancouver, Canada, 28 April – 2 May 2019.

## Figure Legends

**Supplemental Figure 1:**
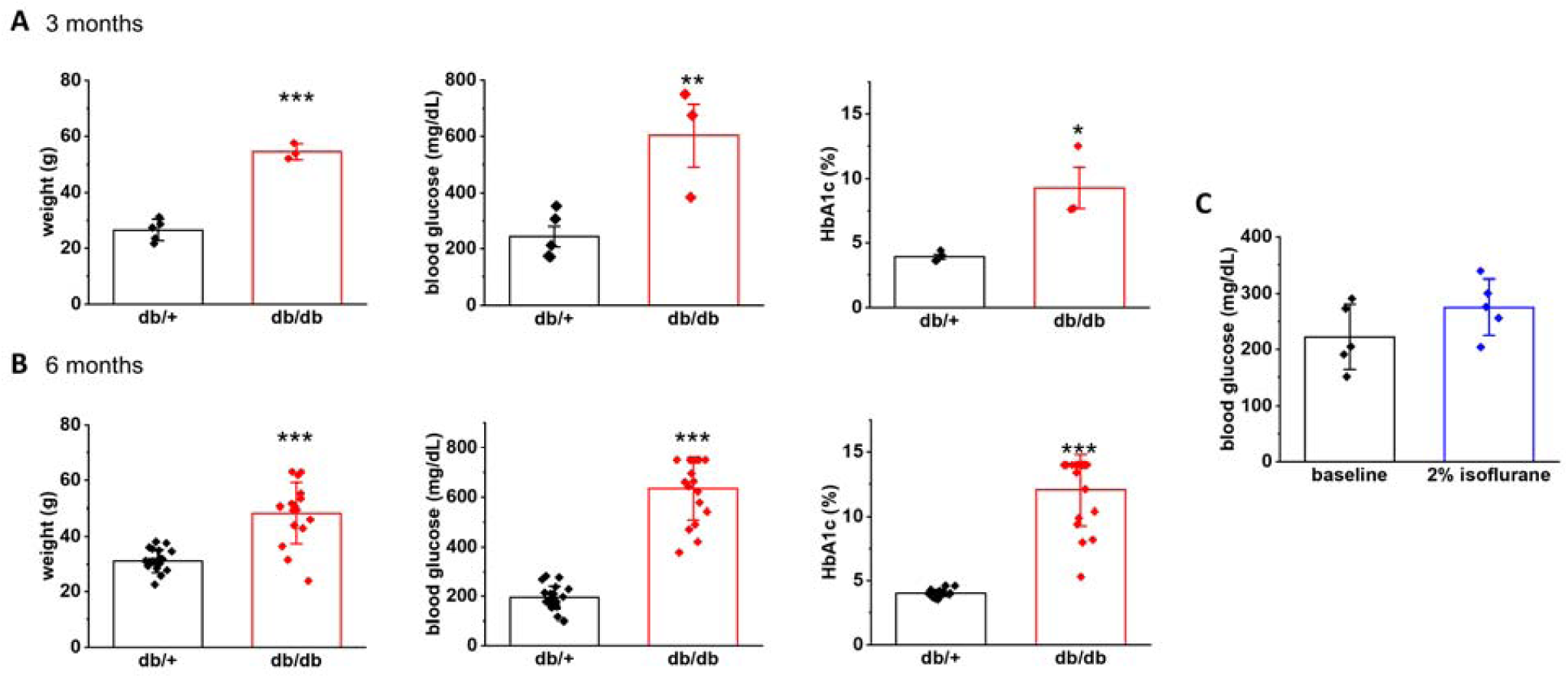
Weight, blood glucose and HbA1c are increased in diabetic mice. Body weight, blood glucose and HbA1c in db/db mice compared to db/+ mice at 3 months (A, n=3-5, * p<0.05, ** p<0.01, *** p<0.001) and 6 months of age (B, n=16-21, ***p<0.001). Blood glucose in C57BL/6J mice before and during 2% isoflurane inhalation (C, n=5)

